# AnNoBrainer, an Automated Annotation of Mouse Brain Images using Deep Learning

**DOI:** 10.1101/2024.01.12.575415

**Authors:** Roman Peter, Petr Hrobar, Josef Navratil, Martin Vagenknecht, Jindrich Soukup, Keiko Tsuji, Nestor X. Barrezueta, Anna C. Stoll, Renee C. Gentzel, Jonathan A. Sugam, Jacob Marcus, Danny A. Bitton

## Abstract

Annotation of multiple regions of interest across the whole mouse brain is an indispensable process for quantitative evaluation of a multitude of study endpoints in neuroscience digital pathology. Prior experience and domain expert knowledge are the key aspects for image annotation quality and consistency. At present, image annotation is often achieved manually by certified pathologists or trained technicians, limiting the total throughput of studies performed at neuroscience digital pathology labs. It may also mean that less rigorous, less time-consuming methods of histopathological assessment are employed by non-pathologists, especially for early discovery and preclinical studies. To address these limitations and to meet the growing demand for image analysis in a pharmaceutical setting, we developed AnNoBrainer, an open-source software tool that leverages deep learning, image registration, and standard cortical brain templates to automatically annotate individual brain regions on 2D pathology slides. Application of AnNoBrainer to a published set of pathology slides from transgenic mice models of synucleinopathy revealed comparable accuracy, increased reproducibility, and a significant reduction (∼50%) in time spent on brain annotation, quality control and labelling compared to trained scientists in pathology. Taken together, AnNoBrainer offers a rapid, accurate, and reproducible automated annotation of mouse brain images that largely meets the experts’ histopathological assessment standards (>85% of cases) and enables high-throughput image analysis workflows in digital pathology labs.

## Introduction

In neuroscience and the study of neurodegenerative disorders, histological methods are widely used to evaluate the pathology present in brain samples of preclinical rodent models of disease [1]. To provide an impartial assessment of the histopathological aspects of a given preclinical model, evaluation should be as unbiased, non-subjective and reproducible as possible. Consistency and quantitative measurements are therefore essential for modeling the pathology present in brain regions at defined stages of the disease, as well as for monitoring the progression of disease over time [2]. Moreover, quantitative evaluation of histological tissue biomarkers, such as markers of cell death [3], pathological protein accumulation [4], or markers of inflammation [5], are similarly critical for successful assessments.

Historically, skilled pathologists or trained technicians have been employed to manually identify specific regions of interest (ROI) in rodent brain samples and to quantify biomarkers of interest across multiple brain regions [6]. Such manual examination is laborious and subjective, limiting the throughput of the development of preclinical rodent models and the testing of therapeutic targets for neurological disorders. In cases where multiple types of image segmentations and annotations are additionally required, the labor and time taken for the entire analysis may increase even further. Moreover, inherent human errors can be amplified by the rote nature of this manual work. A notable example is when pathologists are primed for abnormalities, leading to a bias for false positives.

Opportunities to improve and automate pathological analysis workflows have emerged due to the growing efforts of digitization of histological images and creation of publicly available digital mouse brain atlases. Automated image annotation software tools have the potential to enhance throughput of preclinical histopathological assays, as well as to decrease the subjective evaluation and inherent variability in traditional pathological analysis [7]. Such annotation tools may offer automation of the pathologists’ routine, and consequently free more time for data analysis.

At present there are no commercial tools that provide an automated segmentation of histological rodent brain sections. A few academic tools have been developed and published to date, [8, 9, 10], yet they are limited in terms of the types of Whole-Slide histology Images (WSIs) and histological stains they support. As described by Xu *et al*., existing segmentation models of brain regions are typically based on in-life imaging modalities such as Magnetic Resonance Imaging (MRI) and Computed Tomography (CT) scans, which are at lower resolutions than scanned WSIs. The high-resolution WSIs have discontinuous levels of intensities, preventing the use of segmentation algorithms that rely on the use of homogeneous intensities to identify regions.

In addition, segmentation tools that have been developed for non-neuronal tissues or to differentiate tumor from normal tissues have primarily relied on classification of cell subtypes using machine or deep learning. Such an approach for segmentation of neuronal tissues can successfully detect differences between white and grey matter regions of the brain, but faces greater challenges in separating similar, but spatially differentiated regions, such as differences between cortical regions [11]. Furthermore, most tools that do exist have been trained to focus on a single type of staining, such as Nissl-stained tissue sections or hematoxylin and eosin (H&E) stains, while only a few can accommodate chromogenic and fluorescent immunohistochemical (IHC) assays.

Here we present AnNoBrainer, a novel pipeline for an automated annotation of mouse brain digital pathology images that supports multiple types of Whole-Slide histological Images (WSIs) and histological stains. AnNoBrainer combines murine brain detection using mask-R-CNN (mask region-based Convolutional Neural Network) and an automated brain labelling with user-defined experimental metadata. Moreover, the pipeline incorporates a deep learning classifier for optimal matching of detected brains with brain atlas layers. Furthermore, AnNoBrainer enables accurate non-linear registration enhanced by landmarks-based regularization. AnNoBrainer offloads Pathologists’ repetitive work to a machine and approximately halves the time spent on image annotation while maintaining manual annotation standards. Finally, AnNoBrainer not only streamlines image analysis workflows, but also increases the throughput of digital pathology studies in a pharmaceutical setting. AnNoBrainer is freely available at https://github.com/Merck/AnNoBrainer.

## Methodology

### Digital pathology slides

Data from a study that characterized the aSyn pre-formed fibrils surgical model of synucleinopathy in A30P transgenic mice were used for model validation. Image selection was largely dependent on stain quality and tissue integrity. Significant tears and deformation of the tissue that can dramatically affect detection of morphological boundaries were excluded. Likewise, images were excluded where stain quality was not sufficient for detection of morphological boundaries, or it was substantially different from the staining intensity of neighboring sections. The initial dataset consisted of 313 brains on 19 slides. AnNoBrainer was tested eventually on only 229 brains, since some brains did not pass the initial quality control e.g., due to processing artifacts (e.g., missing half of a brain). Selected brains on each slide were annotated by expert neuroscientists and were limited to three different brain regions as follows: CP (caudioputamen), SNr (substantia nigra) and PAG+AQ (periaqueductal gray + cerebral aqueduct). The final dataset consisted of 89 brains from 17 different layers for CP, 105 brains from 18 different layers for SNr and 35 brains from 7 different layers for PAG+AQ. The distribution of brains in each layer within each region is shown on Supplementary Figure S1.

### Manual annotations

Manual annotations of the caudoputamen (CP) had been made based on morphological boundaries. The lateral edge of the CP was delineated by the corpus callosum and the medial edge by the lateral ventricle. The dorsal CP was separated from the ventral CP by drawing a horizontal line from the most ventral aspect of the lateral ventrical, where it often meets the anterior commissure, across to the lateral edge of the CP. The full striatal area was not captured manually for two reasons. First, the site of injection into the CP only targeted the dorsal portion of the CP. Secondly, accurately, and quickly discerning the ventral CP from the nucleus accumbens using a hematoxylin stain is more challenging than identifying the lateral and medial boundaries of the CP. Using only well-defined morphological markers for manual annotations enabled consistency and minimized biases in drawing the region of interest. However, this also illustrates why manual annotations often sacrifice capturing the full anatomical region of interest in favor of accommodating technical ease.

### Allen mouse brain atlas

The Allen mouse brain atlas [12] consists of 132 coronal sections evenly spaced at 100 μm intervals and annotated to detail numerous brain regions. It was designed to be easily integrated into digital applications in the field of automated histological segmentation [9], and it is considered reliable source of mouse brain anatomy delineating regions of interest, which are mapped and labeled.

### Brain Detection

Detection of brains and hand-written notes on a slide is considered an object detection problem, which consists of two subproblems: 1. detecting the objects of interest (brains and handwritten notes) and 2. segmenting the image with bounding box detection to label the category. For this purpose, a transfer learning strategy [13] was followed, and a pre-trained MASK R-CNN model [14] was modified by replacing its last fully connected layer with a set of new fully-connected layers retrained on 20 manually labeled slides aiming to detect two distinct categories – 1) mouse brains and 2) handwritten notes. For the training process, ADAM optimizer was used with 60 epochs and batch size of 2. Following brain detection, bounding boxes were generated for each individual mouse brain on a slide and centroids were calculated to form a brain centroid grid.

### Linking of Detected Images

To differentiate between experimental and control group brains, an experimental table was manually created for each slide. To link the detected brains from the slides to their corresponding descriptions in the experimental table, the following approach was employed. First, a brain centroid grid from the brain detection step was obtained. A similar grid was then constructed from the experimental table using its row and column coordinate structure. Both grids were then standardized to ensure that they were in the same unit of measurement. To connect the corresponding points on each grid, Hungarian algorithm [15] was applied. The algorithm assigns each pair from both tables in such a way that the total sum of distances is minimized.

### Matching with Reference Atlas Layer

Matching of brains to their respective reference (z-slice) layers is a manual, non-trivial and subjective task that requires expert knowledge. Multiple convolution neural networks architectures for classification were trained and tested, using a standard test/train split approach and image augmentations, including rotation, random brightness, channel flip, median blur, and elastic transforms [16]. The latter is an important component of the training since individual brains are cut by hand and a certain level of asymmetry is generally present. The ResNet34 [17], and Efficient Net architectures were used [18]. Since this task can be also interpreted as a regression problem, a Label Smoothing Cross-Entropy Loss function [19] were used for the training. The training used pre-trained models on the ImageNet dataset and only replaced the last fully connected layers that are responsible for the classification itself. These were fully trained from scratch using a batch size of 12 and ADAM optimizer for 12 epochs.

### Image Registration Process

Image registration is an iterative optimization process that searches for best suitable geometrical transformation to spatially align two images, a reference image, and a target image. The image registration process implemented in our pipeline consists of subsequent steps of geometric alignment between reference Allen mouse brain atlas layer and mouse brain identified on a slide.

Airlab Library [20] and the detection of sparse cross-domain correspondences [21] were the main tools utilized for AnNoBrainer’s custom registration pipeline. AirLab provides flexibility with various image registration methods, loss functions, and regularization techniques all using Pytorch, and can be run on a GPU cluster. Affine Registration was the first step in our procedure followed by Elastic Registration to correct local nonlinear disparities. The objective function for image registration was extended by considering a sparse correspondence detected in target image and in the template image via [21]. Thereafter, the elastic registration regularization term was extended to minimize the distance between the corresponding landmarks while ensuring smooth geometric transformation.

### Affine Registration

Affine alignment is a technique used to align two images by applying 2D linear image transformations and translations. The transformations include operations such as scaling, reflection and rotation, etc. The goal is to minimize the loss function by gradually transforming the registered image to make it as similar to the template image as possible. Similarity transformation was used to preserve the original shape of the image, which was transformed by simple operations such as rotation and reflection. The Normalized Cross-Correlation loss objective function was used for the affine registration process. To perform the registration, Airlab’s backend was used to allow iterative GPU-based optimization. The optimization software gradually transformed the registered image to make it as similar to the template image as possible. The ADAM optimizer was used for the optimization process with a learning rate of 0.01 and 1000 iterations.

### Elastic Registration

Once the affine registration aligns the image to a template image, elastic registration follows. Elastic registration estimates a smooth continuous function to map the points in a registered image to a template image. Three types of transformation models are common: linear/dense, non-linear/interpolating, and dense models. AnNoBrainer employed the non-linear/interpolating approach. To transform point *x* in the image, the displacement *f*(*x*) is interpolated from neighboring control points. The interpolation is done using a basis function. For the basis function, a B-spline kernel was used. A diffeomorphic approach was employed to ensure the preservation of topology. The affine registration was extended with a regularization term. To limit the number of sampling points, a Regulariser (DiffusionRegulariser) that penalized changes in the transformation was used on the displacement field. Like the Affine registration, the normalized cross-correlation loss function was used for optimization. Adam optimizer with a learning rate of 0.01 and 1000 iterations was used to optimize the transformation field.

### Custom Landmark Regularization Term

During the registration process, two common edge cases occur, the distortion is either too high or too low. This is usually a result of a balance between the loss function and regularization term. To achieve realistic and anatomically reasonable distortions, a higher regularization and smaller distortions approach is typically used. A common issue encountered is when the tissue cut is not perpendicular to the z-axis of the brain. This results in an anatomical difference between both hemispheres of the brain, causing imperfect alignment in some parts of the brain when relatively high regularization is used. To overcome this, a higher level of flexibility is required. Therefore, an additional step was implemented to higher or lower regularization term for certain parts of the image and allow higher distortion locally. Image landmarks serve as the layer of the pipeline that fills the gap. A landmark in image processing may be understood as a characteristic feature or structure in any given image. Landmarks across two images are points at the same semantic locations, e.g., left edge of the organ. When Landmarks are aligned, the regularization term is the same as it would be without landmarks. However, when there is a significant difference between landmarks (landmarks of template image and landmarks of registered image) it locally forces regularization term to have less effect. This leads to (possibly) overall better image alignment. To find landmark points between two images, a pre-trained neural network from Neural Best-Buddies: Sparse Cross-Domain Correspondence [21] was used to identify semantically matching areas for two cross-domain image pairs. Then a landmarks-derived regularization term was constructed. The aim was to minimize the squared Euclidean distance between corresponding landmarks. The squared Euclidean distance was preferred on the classic square root version, given that without the square root the smoothness of the function at all points is preserved, which is an important aspect for gradient optimization.

A Landmark regularization term (*LRT*):

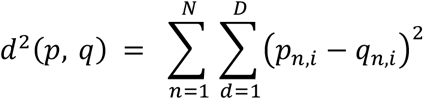

Where *p* denotes warped landmark point from moving image, *q* denotes landmark point from fixed image, *N* represents number of landmark point and *D* represents a dimension corresponding to x and y position in image.

An elastic Regularization loss uses Normalized Cross Correlation (*NCC*) extended with a Diffusion Regulariser term (*DF*). Thus, the loss function extended with a Landmark regularization term is defined as follows:

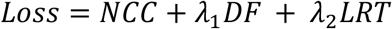

Where terms *λ*_1_ and *λ*_2_ are both selected as 0.5.

## Results

### AnNoBrainer features seamless mouse brain annotation using deep learning

Histopathological assessment of preclinical murine models is instrumental for neurodegenerative studies, yet it is largely dependent on manual image analysis and annotation by experienced pathologists, which is time-consuming, subjective, not consistent, or scalable. To fill in this gap AnNoBrainer offers an end-to-end pipeline that significantly accelerates digital pathology campaigns. The pipeline is comprised of four main steps (Figure 1), as follows. First, the tool detects individual brain sections present on the 2D slides using mask-RCNN deep learning model that also efficiently identifies and disregards noise on the physical slides such as hand-written notes (Figure 1A). Second, using Hungarian algorithm [15] AnNoBrainer automatically labels individual brains with user-defined experimental metadata provided in an Excel spreadsheet format (Figure 1B). Thereafter, a deep learning classifier is employed to identify the optimal Allen mouse brain atlas template for each brain (Figure 1C). Once the reference layers for all individual brains are identified, image registration for each image pair is taking place to determine optimal transform parameters (1D). Subsequently, user-selected ROIs are transferred from the reference atlas layer to the target brains ROIs (1E).

**Figure 1.**
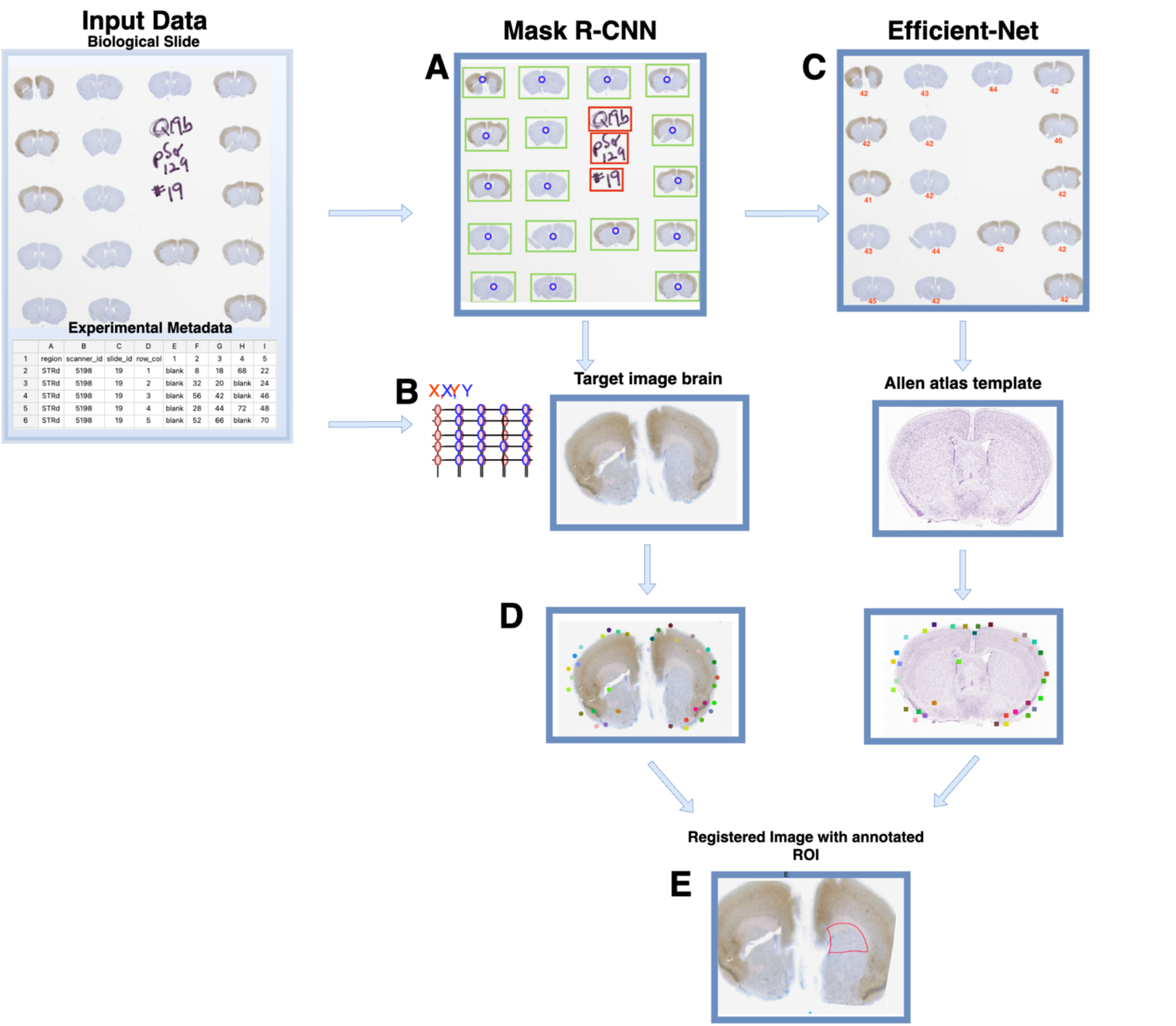
AnNoBrainer enables seamless and automated identification of brain objects from multi-brain slides, their matching to experimental metadata and reference brain templates as well as efficient and robust image registration. (A) a mask-RCNN identifies brains and background handwritten notes (B) an Hungarian algorithm is used to link experimental metadata with individual brains (C) Efficient-Net model is used for matching a target brain with its corresponding Allen atlas layer (D) image registration of brain with Allen atlas template using affine and elastic registration that is enhanced by custom landmark regularization (E) region of interest is geometrically transformed to target brain.

### AnNoBrainer exhibits flawless detection of brain and background objects

Often, in a digital pathology laboratory, murine brain sections from multiple individuals are placed on a single slide and typically accompanied by hand-written notes. Therefore, the annotation pipeline must identify and distinguish between brains and background objects. For this task, AnNoBrainer employs a pre-trained mask R-CNN model (Table 1) that was further extended to include two designated ‘brain’ and ‘note’ categories. The model was then fine-tuned to distinguish between these two new classes. Briefly, for training purposes 20 slides were manually annotated with bounding boxes representing the respective two new classes. To ensure that the model will be able to generalize effectively in the future, the slides for the training were selected in such a way that they were highly variable and represented a broad spectrum of use cases, as follows. Labelled slides accounted for instances with randomly missing brains, rotated and torn tissue, and instances with many or no hand-written notes at all.

**Table 1.** Deep learning models used in this study.

| Model Type | Function | Input | Output |
| --- | --- | --- | --- |
| Mask R-CNN | Detects Individual Brains and handwritten notes present on the slide | Slide image | Coordinates of individual brains present on the slide |
| EfficientNet-B0 | Match brain to the most similar Allen brain atlas reference layer | Individual brain image | Predicted Z-axis from the Allen brain atlas |

The fine-tuned mask R-CNN model was then applied to 20 new slides, where it perfectly identified and segmented all individual brains on each of the slides as well as correctly classified all hand-written note objects.

### AnNoBrainer efficiently links detected brain images to experimental metadata

Experimental metadata associated with the brain images are critical for downstream data analysis. AnNoBrainer enables seamless matching between user provided metadata (Excel table format) and their respective brain images. Briefly, the Excel table containing the metadata is first converted to a coordinate grid (Figure 1B, red). Once individual brains are detected on a slide, their centroids are computed to derive a second grid (Figure 1B, blue). Both grids are then aligned via standardization. Euclidean distance between all points in both grids is computed and Hungarian algorithm is employed to assign each brain with its associated unit grid node.

### AnNoBrainer matches brain images to reference brain atlas templates with high accuracy

Correct classification of brain layers is crucial for experimental neuroscience given the spatial gene expression patterns across different layers and cell types. Databases such as the Allen mouse Brain Atlas provide a comprehensive anatomical brain map that can be integrated into histopathological analysis workflows. Although matching brain layers to detected brain objects can be performed manually by an expert pathologist, it is not a trivial task and is somewhat subjective. AnNoBrainer offers expert users to either manually select the corresponding reference brain template for each brain image or alternatively to use an automated matching approach. For the latter, it employs a deep learning model trained on labelled data that were manually curated by expert pathologists. To ensure variability within the training set, images were selected in all shapes and forms, including clean and symmetrical brains alongside asymmetrical and slightly torn brain sections. Similarly, images used for training were resized to 460 pixels to ensure consistency. Importantly, training data only accounted for a subset of atlas z-slices (39-73 CP, Figure S2), each containing expert-labelled images, limiting the prediction of templates to this spectrum of z-slices. Selected examples of labelled images for reference brain atlas training are shown in Figure S3. Multiple Convolutional Neural Network (CNN) were initially tested using standard data train-test split and image augmentation (e.g., brightness, blur, rotation etc.,) fine-tuned to the extent that training and testing losses ceased to decrease any further. The EfficientNet-B0 model outperformed the ResNet-34 model at default or fine-tuned states, reaching an overall 94% accuracy level when a misclassification tolerance of 1-2 neighboring layers with respect to expert annotation was applied (Tables 1 and 2).

### AnNoBrainer offers a robust image registration technique that ensures optimal spatial alignment

Once a 2D reference brain template is identified it needs to be spatially aligned to its respective physical brain image on the slide via an iterative optimization process known as image registration. AnNoBrainer employs robust, non-linear image registration techniques that combines Affine and Elastic registration approaches for geometric image alignment and smoothing, respectively. To further improve image alignment, AnNoBrainer incorporates an optional automated detection of semantic locations on the images that essentially act as anchor characteristic features between the image pair, also referred to as custom landmark terms (Figure 1D).

**Table 2.**
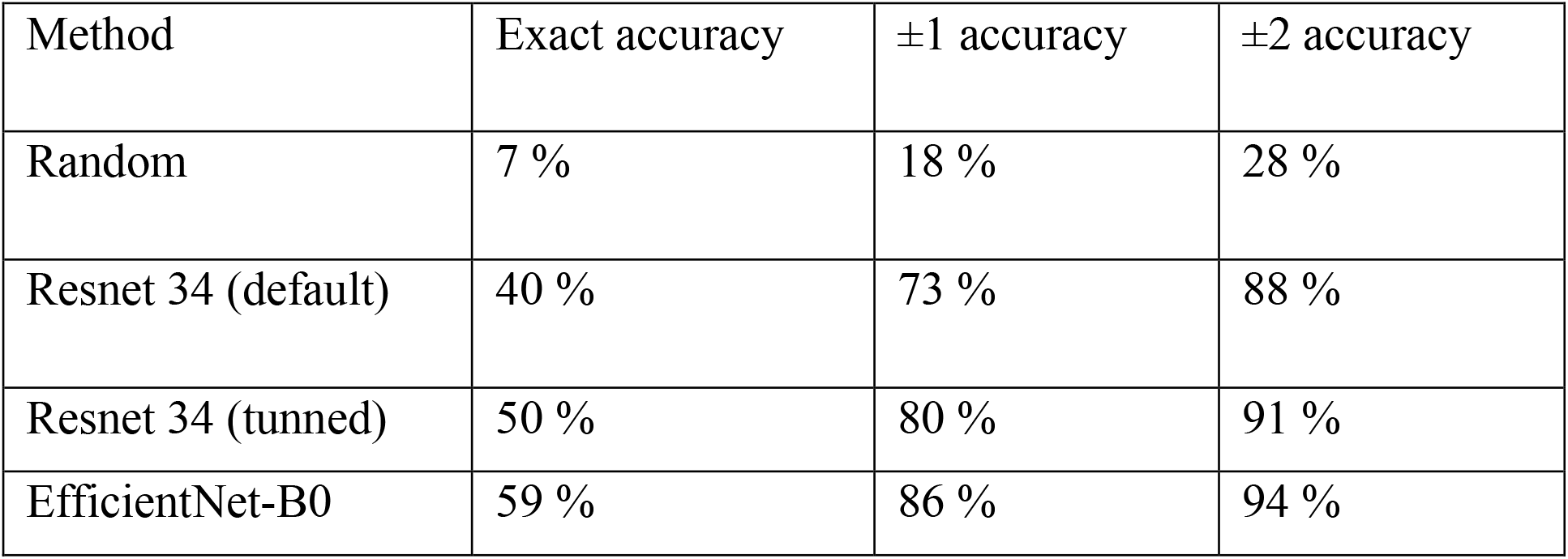
Model performances for Brain atlas layers matching.

| Method | Exact accuracy | $\pm 1$ accuracy | $\pm 2$ accuracy |
| --- | --- | --- | --- |
| Random | 7 % | 18 % | 28 % |
| Resnet 34 (default) | 40 % | 73 % | 88 % |
| Resnet 34 (tunned) | 50 % | 80 % | 91 % |
| EfficientNet-B0 | 59 % | 86 % | 94 % |

The performance of the automated registration was evaluated using a segmentation metric between the expert annotated ROIs and the ROIs automatically registered by AnNoBrainer. In a semantic segmentation the aim is to predict a mask. Put differently, to predict where the actual object of interest is present. Here, both the expert annotated, and the automatically registered ROIs were treated as a binary mask i.e. 0 and 1. Thereafter, F1 score was used to quantitatively assess the registration accuracy by measuring the similarity between expert-annotated and predicted ROIs for each image pair. Furthermore, the effect of custom landmarks terms on registration quality was also evaluated.

Regardless of the brain region tested (Caudoputamen - CP, Periaqueductal gray - PAG, and Substantia nigra - SNr) image registration quality was significantly improved when AnNoBrainer automated atlas layer detection was combined with custom landmark terms, compared to manual layer selection by expert without the use of landmark terms (Figure 2A-B, t-test p <0.00013).

**Figure 2.**
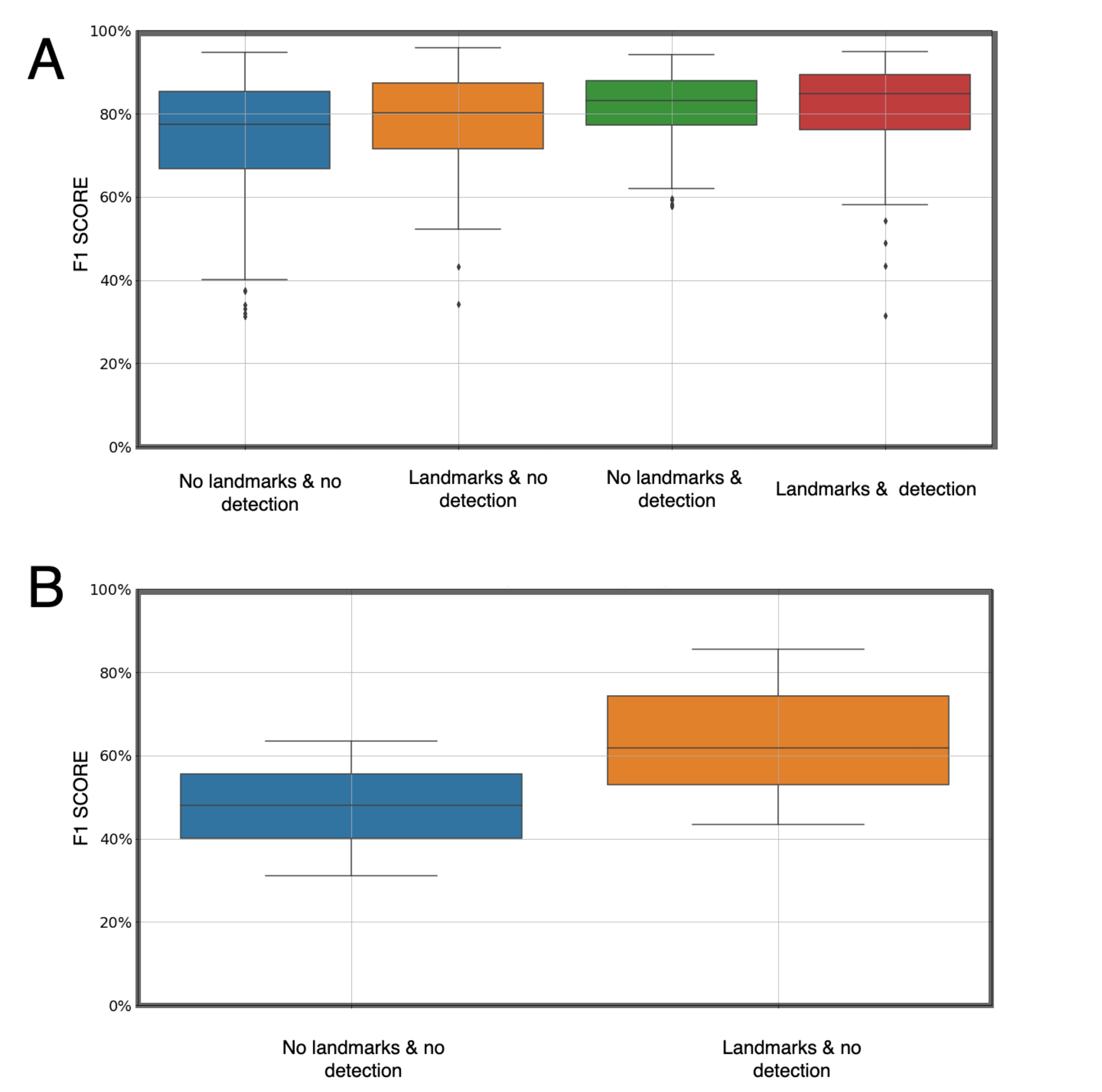
Registration accuracy for CP and PAG regions respectively. (A) registration accuracy for CP region with various pipeline settings, as indicated on the x-axis; (B) registration accuracy for PAG region with various pipeline settings, as indicated on the x-axis

Finally, a comparison between manual annotation and expert annotation was performed revealing that the majority (>67%) of AnNoBrainer automated annotations met the expert annotation standards while some (∼18%) needed minor adjustments (Figure 3 A-B). In approximately 15% of the cases AnNoBrainer completely failed to precisely annotate the brain images within accepted boundaries, primarily due to discrepancies in structural information between the input data and the Allen brain atlas (Figure 3C).

**Figure 3.**
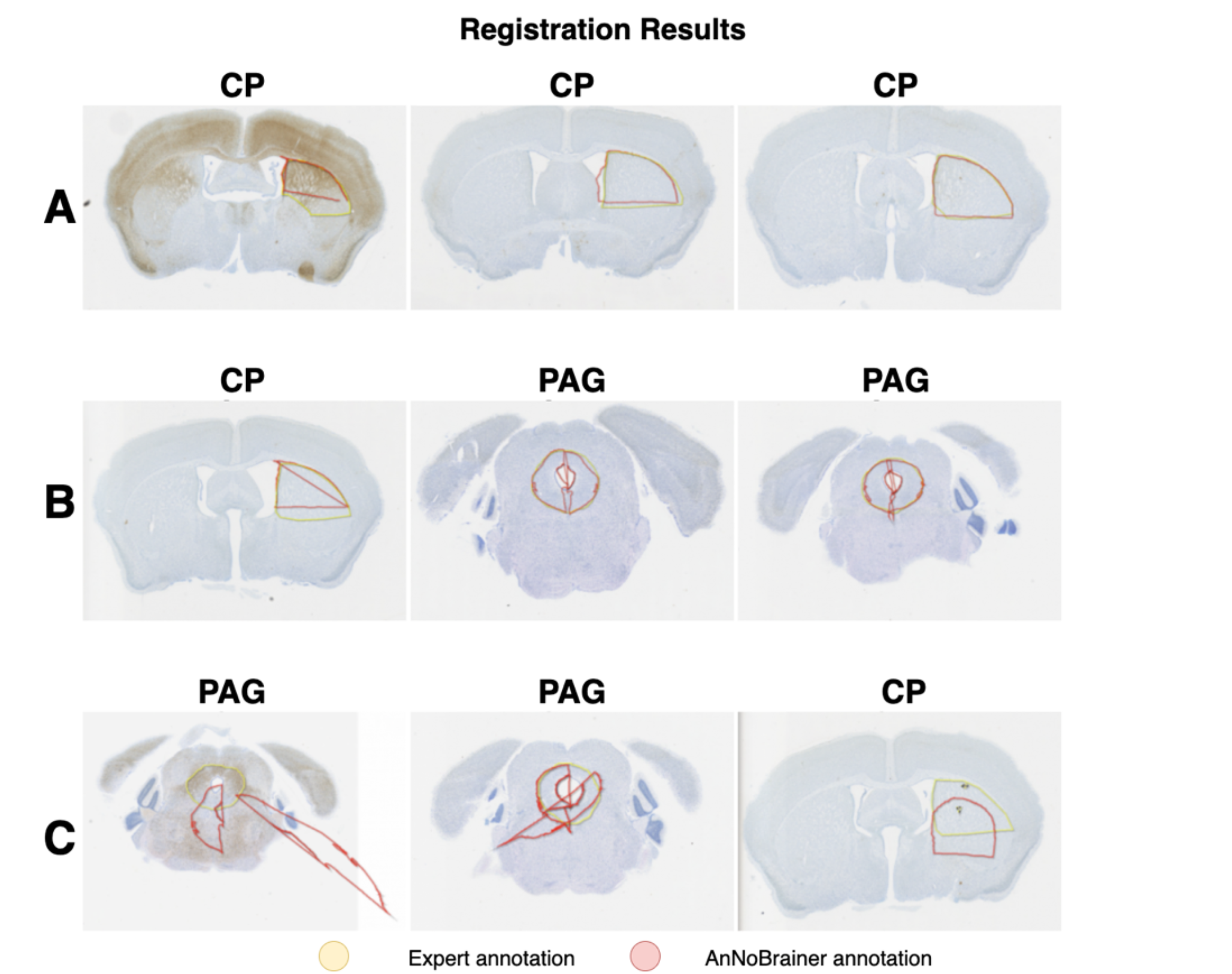
Registered ROI’s by AnNoBrainer. (A) Satisfactory annotation not requiring manual adjustments, (B) Satisfactory annotation requiring slight manual adjustments, (C) Unsatisfactory annotation requiring significant manual adjustments

## Discussion

Pre-clinical neuroscience research relies largely on histopathological analysis of brain samples of rodent models of disease. The emergence of digitized pathology slides and advanced machine and deep learning methods opens new opportunities to automate, streamline and scale up laborious analysis workflows without compromising the quality achieved by human experts. AnNoBrainer has been successfully applied in multiple high-throughput pre-clinical neuroscience studies in the industry, and it performed well on a large number of slides with a varying quality of brains (e.g., slightly torn, asymmetrical etc.), different stains (e.g. H&E, Nissil, IHC), background noise and various templates. Apart from the benefit of automation of mundane work, AnNoBrainer also reduces the annotation time of a given slide by approximately 50%.

Despite these advantages it is also important to note AnNoBrainer’s limitations. The atlas layer identification model in its current form is only applicable to a limited number of brain slides (z-direction), since it was trained only on slides within the CP region. This limitation can be readily addressed by retraining the model with an extended labelled dataset. Another caveat is that the image registration quality of AnNoBrainer may be affected by the quality of the input brains with respect to torn or missing tissue, where asymmetry or tears found in close proximity to the registered region may result in poor registration around these areas. Additional drawbacks that are associated with the matching of brain layer template and the registration process can be attributed to the discrepancies between the varying layer sizes in the Allen brain atlas and the actual size of the brain slices provided as input. Similarly, certain IHC staining techniques such as DAPI are unlikely to result in high quality registration due to data sparsity or poor morphological correspondence with the H&E staining used in the Allen brain atlas. Important to note that to date Nissl, H&E and some IHC stains (e.g. α-synuclein) were successfully tested, yet AnNoBrainer was designed to also support multiplexed stains and can incorporate other IHC stains too, therefore can support a broader range of studies compared to previously published methods [11, 10] that focus primarily on traditional histology stains. L

## Conclusion

In this study we developed an automated, deep learning-based pipeline for mouse brain annotation. Our pipeline can efficiently identify and distinguish mouse brain from noise, link it to its respective experimental metadata and register it with its corresponding brain layer template. AnNobrainer was designed as a modular and extensible annotation pipeline to allow retraining of models and/or introduce further improvements by the community as more labelled data become available or when other staining techniques are preferred.

## Availability

### Availability of data and materials

All code for this publication is available in the following GitHub repository: https://github.com/Merck/AnNoBrainer.

## Competing Interests

All authors that are/were employees of Merck Sharp & Dohme LLC, a subsidiary of Merck & Co., Inc., Rahway, NJ, USA and may hold stocks and/or stock options in Merck & Co., Inc., Rahway, NJ, USA. A subset of manuscript authors are inventors on a patent related to this work (patent application number: 34187-56244). All of this work/code is licensed under the MIT permissive free software license.

## Funding

This work was supported by Merck Sharp & Dohme LLC, a subsidiary of Merck & Co., Inc., Rahway, NJ, USA.

## Author Contributions

RP, DB, JSU, MJ, RG conceived the study. RP supervised the study an orchestrated the development. PH, MV, JS and JN developed and implemented the entire solution. KT, NB and AS tested and validated the models. DB wrote the manuscript. PH, JN, RP, MV, RG and JSU wrote the initial draft. All authors contributed to the final draft. All authors reviewed and approved the final draft.

## Acknowledgments

We thank Jens Christensen, Jyoti Shah, Antong Chen, Ondrej Holub, and Carol A. Rohl for supporting this work. We are immensely grateful to for their advice and stimulating discussions.

## Notes

### Summary of Updates

Author full names

