## Supplementary Figure S1 for "AnNoBrainer, an Automated Annotation of Mouse Brain Images using Deep Learning"

### Supplementary Material

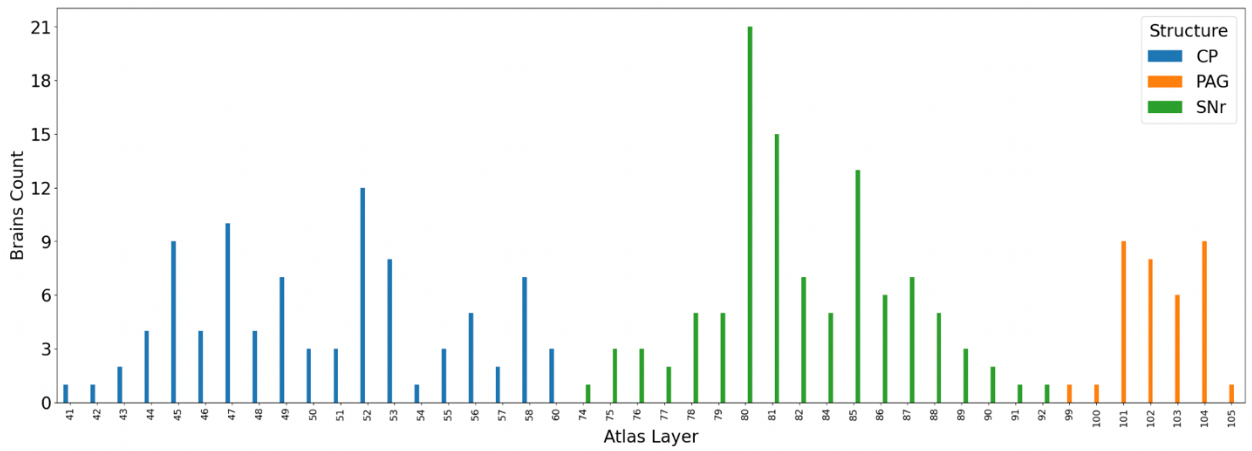


**Figure S1.** Dataset for validation of our pipeline. It shows number of brains in the dataset depending on Atlas Layer number, which is taken from the Allen Mouse Brain Atlas. Different regions in our dataset are distinguished by the color of the bar.

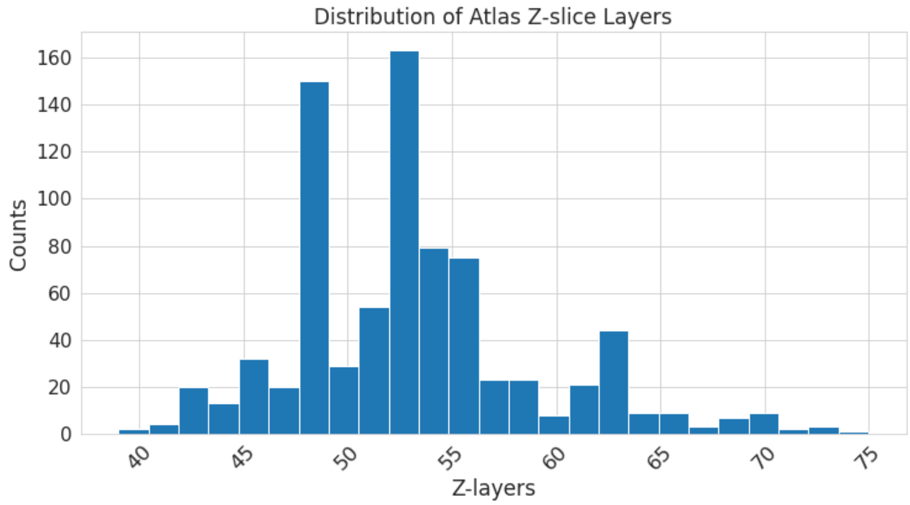


**Figure S2**. Distribution of atlas slices used for training of the brain to layer matching models.

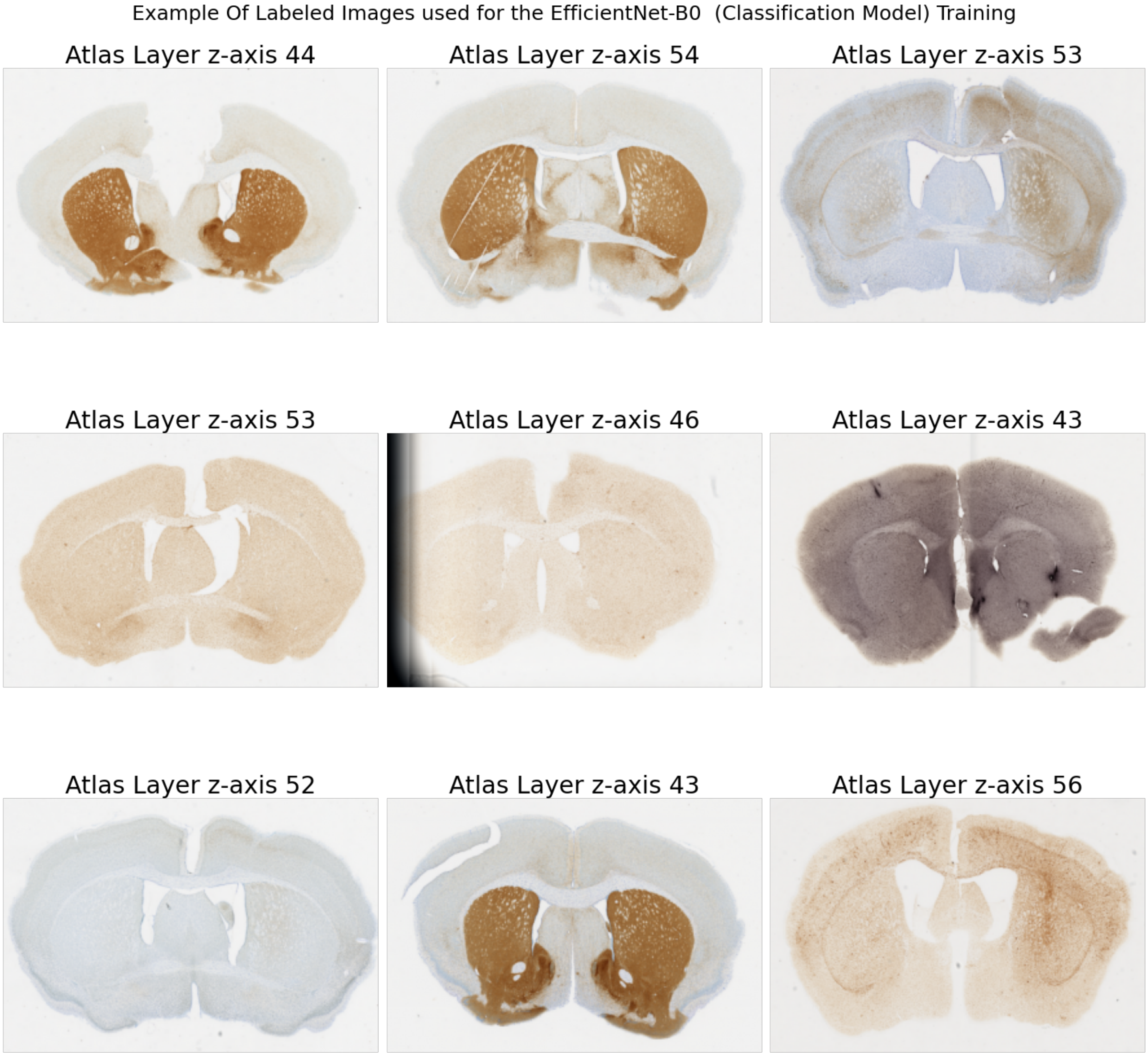


**Figure S3**. Distribution of atlas slices used for training of the brain to layer matching models.
